# Expression of Unfolded Protein Response Genes in Post-transplantation Liver Biopsies

**DOI:** 10.1101/2022.02.21.480553

**Authors:** Xiaoying Liu, Sarah A. Taylor, Stela Celaj, Josh Levitsky, Richard M. Green

## Abstract

Cholestatic liver diseases are a major source of morbidity and mortality that can progress to end-stage liver disease. There are few effective medical therapies for primary biliary cholangitis, primary sclerosing cholangitis and other cholestatic liver diseases, in part, due to our incomplete understanding of the pathogenesis of cholestatic liver injury. The hepatic unfolded protein response (UPR) is an adaptive cellular response to endoplasmic reticulum stress that is important in the pathogenesis of many liver diseases and recent animal studies have demonstrated the importance of the UPR in the pathogenesis of cholestatic liver injury. However, the role of the UPR in human cholestatic liver diseases is largely unknown. In this study, we utilized liver biopsies from patients after liver transplantation as a disease model to determine the transcriptional profile and hepatic UPR gene expression that is associated with liver injury and cholestasis. RNA-seq analysis revealed that patients with hyperbilirubinemia had enhanced expression of hepatic UPR pathways. Alternatively, liver biopsy samples from patients with acute rejection had enhanced gene expression of *LAG3* and *CDK1*. Pearson correlation analysis of serum alanine aminotransferase, aspartate aminotransferase and total bilirubin levels demonstrated significant correlations with the hepatic expression of several UPR genes, as well as genes involved in hepatic bile acid metabolism and inflammation. In contrast, serum alkaline phosphatase levels were correlated with the level of hepatic bile acid metabolism gene expression but not liver UPR gene expression.

**Conclusion:** Overall, these data indicate that hepatic UPR pathways are increased in cholestatic human liver biopsy samples and supports an important role of the UPR in the mechanism of human cholestatic liver injury.

---

Cholestatic liver diseases including primary biliary cholangitis (PBC), primary sclerosing cholangitis (PSC), biliary atresia and familial genetic etiologies remain a major source of significant morbidity and mortality. Unfortunately, there are few effective medical therapies for PBC, and no effective medical therapies for PSC and many genetic cholestatic liver disorders, and liver transplantation is the only life-saving option for end-stage diseases (1, 2). In addition, following liver transplantation, patients with certain cholestatic liver diseases can have significant post-transplantation recurrence rates, with rates of up to 53% in PBC and up to 45% in PSC (2-6). Post-transplantation liver disease recurrence may result in patient graft loss or death. A major reason for the lack of effective medical therapies for cholestatic liver disorders is our incomplete understanding of the disease pathogenesis and progression. Recent human and animal studies indicate that the liver unfolded protein response (UPR) is important in the pathogenesis of cholestatic liver injury and may be prognostic for liver-related complications in patients with PSC (7-9).

The UPR is an adaptive cellular response to endoplasmic reticulum (ER) stress. ER stress is a form of cellular stress that occurs due to an accumulation of excess unfolded or misfolded proteins in the ER. Since protein synthesis in the liver is quantitatively high, it may be particularly susceptible to the development of ER stress (10, 11). The UPR functions to reduce the number of cellular misfolded or unfolded proteins by enhancing protein folding, attenuating protein translation, and increasing endoplasmic reticulum-associated protein degradation. However, if ER stress is severe and cannot be resolved, it activates apoptosis pathways. The UPR is comprised of three signaling pathways including inositol requiring enzyme 1α/X-box binding protein 1 (XBP1), PKR-like ER kinase (PERK) and activating transcription factor 6 (ATF6), that regulate downstream UPR genes to return cellular homeostasis (12, 13). The hepatic UPR is important in the pathogenesis of many liver diseases including viral hepatitis, non-alcoholic fatty liver disease, alpha-1 antitrypsin deficiency, alcoholic liver disease and ischemia-reperfusion injury (9-11). Finally, in a recent study of PSC patients, differential expression of UPR genes was identified in patients who were at high risk for liver-related complications (7). Unfortunately, the role of the UPR in human cholestasis and cholestatic liver injury remains poorly understood. In order to better determine the role of the hepatic UPR in the pathogenesis of human liver disease, we performed transcriptome analysis on “for-cause” (clinically-indicated for graft injury/dysfunction) liver biopsies from post-transplantation patients and sought to determine how changes in hepatic UPR gene may be associated with liver injury and cholestasis.

## Methods

### Human samples

Twenty liver transplant recipients (2013-2015) undergoing a for-cause liver biopsy at Northwestern Memorial Hospital consented to have a portion of their liver biopsy utilized for this study. Briefly, liver biopsy was performed with a 16 gauge 33 mm BioPince needle. If adequate sample size was obtained (>2 cm) for routine histology, a 0.5-1 cm piece was removed from the end of the main piece, placed in RNAlater and stored at - 80°C. Patient demographics, laboratory tests, medication, and clinical data were collected and utilized from the Northwestern Medicine ® Enterprise Data Warehouse, which is a single, comprehensive and integrated repository of clinical and research data sources. The biopsies were locally reviewed for clinical care purposes and then also underwent an independent, blinded central review. Acute rejection (AR) was scored using the Banff Rejection Activity Index (14). Clinical and histological data were reviewed by a transplant hepatologist (J.L.). Liver biopsies were categorized as: 1) *AR* if histology demonstrated evidence of acute rejection; 2) *Non-Rejection: hyperbilirubinemia (NR:HBR)* if serum total bilirubin was > 2.5 mg/dL and there was no histologic evidence of rejection; 3) *NR: normal or mild elevation in liver function tests (LFTs) (NR:Mild)* if there was no histologic evidence of rejection, serum total bilirubin ≤ 2.5 mg/dL and alanine transaminase (ALT), aspartate aminotransferase (AST) and alkaline phosphatase (ALP) levels were ≤ 1.67x upper limit of normal (15); and 4) *NR: others with non-HBR high LFTs (NR:Others)* if there was no histologic evidence of acute rejection, but serum ALT, AST and ALP levels were > 1.67x upper limit of normal with total bilirubin ≤ 2.5 mg/dL. This study was approved by the Northwestern University Institutional Review Board (STU00213022). No donor organs were obtained from executed prisoners or other institutionalized persons.

### RNA-seq analysis

Total RNA was isolated from liver biopsies using the RNeasy micro kit (Qiagen, Germantown, MD) according to the instructions of the manufacturer. RNA-seq was conducted at Northwestern University NUSeq Core Facility as recently described (16). Briefly, total RNA samples were checked for quality using RNA integrity numbers (RINs) generated from the Agilent Bioanalyzer 2100. One sample failed QC (RIN < 7) and was excluded from the study, therefore 19 samples proceeded to sequencing. RNA quantity was determined with Qubit fluorimeter. The Illumina TruSeq Stranded mRNA Library Preparation Kit was used to prepare sequencing libraries from 1μg of high-quality RNA samples (RIN > 7). This procedure includes mRNA purification and fragmentation, cDNA synthesis, 3’ end adenylation, Illumina adapter ligation, library PCR amplification and validation. An lllumina HiSeq 4000 sequencer was used to sequence the libraries with the production of single-end, 50 bp reads at the depth of 20-25 M reads per sample.

The quality of reads, in FASTQ format, was evaluated using FastQC. Reads were trimmed to remove Illumina adapters from the 3’ ends using cutadapt. Trimmed reads were aligned to the human genome (hg38) using STAR (17). Read counts for each gene were calculated using htseq-count in conjunction with a gene annotation file for hg38 obtained from Ensembl (http://useast.ensembl.org/index.html). Normalization and differential expression were calculated using DESeq2 that employs the Wald test (18). The cutoff for determining significantly differentially expressed genes was an FDR-adjusted p-value less than 0.05 using the Benjamini-Hochberg method. Ranking of differentially expressed genes in *NR:Mild* and *NR:HBR* groups was performed using the EdgeR package in R studio version 1.2.1335 (19-21).The normalized enrichment score for hallmark gene sets was then determined using gene set enrichment analysis (GSEA) software (22, 23). In addition, data analysis was also performed using GeneCodis 4.0 to identify significant pathways among the significantly differentially expressed genes (https://genecodis.genyo.es/). A P-adj value <0.05 was deemed to be statistically significant. Lastly, comparisons between serum liver chemistries (ALT, AST, total bilirubin and ALP) and hepatic gene expression data was preformed using Pearson Correlation test in PRISM 9 software (GraphPad, San Diego, CA). Statistical significance was defined as a P-value of less than 0.05.

## Results

Table 1 and Supplementary Table 1 list the patient demographics, clinical information and biopsy category as defined in Methods for nineteen patients. Three patients had evidence of AR, while there was no histologic evidence of AR in the sixteen other liver biopsies. Of the NR groups, 3 patients were categorized as *NR:HBR*, 4 patients were categorized as *NR:Mild*, and 9 patients were categorized as *NR:Others*. The patients in the

**Table 1:** Patient Characteristics.

|  | **AR**<br>**(n=3)** | **NR:HBR**<br>**(n=3)** | **NR:Mild**<br>**(n=4)** | **NR:Others**<br>**(n=9)** |
| --- | --- | --- | --- | --- |
| **Age at Transplant**<br>**(years, mean [range])** | 44 [20, 64] | 45 [27, 65] | 47 [26, 65] | 54 [25, 65] |
| **Caucasian Race (%)** | 3 (100) | 3 (100) | 3 (75) | 7 (78) |
| **Male Sex (%)** | 1 (33) | 2 (67) | 3 (75) | 4 (44) |
| **Primary Liver Diagnosis (%)** |  |  |  |  |
| Hepatitis C (non-viremic) | 0 (0) | 0 (0) | 0 (0) | 1 (11) |
| Alcohol | 1 (33) | 0 (0) | 1 (25) | 0 (0) |
| Non-Alcoholic Fatty Liver or<br>Cryptogenic | 1 (33) | 0 (0) | 1 (25) | 3 (33) |
| Immune-Mediated<br>(PSC, AIH, PBC) | 0 (0) | 3 (100) | 2 (50) | 4 (44) |
| Other | 1 (33) | 0 (0) | 0 (0) | 1 (11) |
| **Months from LT (mean, [range])** | 8.0 [4.6, 10.2] | 44 [0.8, 71.5] | 82 [6.5, 119.1] | 63 [4.7, 142.5] |
| **Immunosuppression (%)** |  |  |  |  |
| CNI therapy | 3 (100) | 3 (100) | 4 (100) | 9 (100) |
| Mycophenolic acid therapy | 3 (100) | 3 (100) | 2 (50) | 2 (22) |
| Predisone | 1 (33) | 3 (100) | 1 (25) | 4 (44) |
| **Laboratory Values**<br>**(mean, [range])** |  |  |  |  |
| ALT (U/L) | 172 [68, 252] | 165 [27, 302] | 54 [20, 84] | 98 [22, 303] |
| AST (U/L) | 115 [27, 178] | 104 [50, 134] | 41 [18, 64] | 44 [22, 103] |
| Alkaline phosphatase (U/L) | 146 [54, 226] | 264 [164, 366] | 100 [68, 130] | 363 [191, 621] |
| Total Bilirubin (mg/dL) | 1 [0.5, 1.2] | 8 [3.5, 12.2] | 1 [0.5, 1.5] | 1 [0.3, 1.5] |
| **Rejection Characteristics (%)** |  |  |  |  |
| Mild (RAI 3-4) | 1 (33%) | -- | -- | -- |
| Moderate - Severe (RAI 5-9) | 2 (67%) | -- | -- | -- |

**AR:** acute rejection
**NR:Mild:** non-rejection; serum total bilirubin ≤ 2.5 mg/dL; ALT, AST and ALP ≤ 1.67x ULN
**NR:Others:** other non-rejection with serum total bilirubin ≤ 2.5 mg/dL; ALT, AST and ALP > 1.67x ULN

### *NR:Mild* and *NR:HBR* groups had serum total bilirubin levels that were ≤ 1.5 mg/dL

Figure 1 depicts the principal component analysis (PCA) of RNA-seq data performed on the 19 samples. PCA analysis demonstrated that the 3 samples in the *NR:HBR* group clustered independently from all other samples. In contrast, samples from the 3 other categories did not cluster independently from any of the other groups.

**Figure 1:**
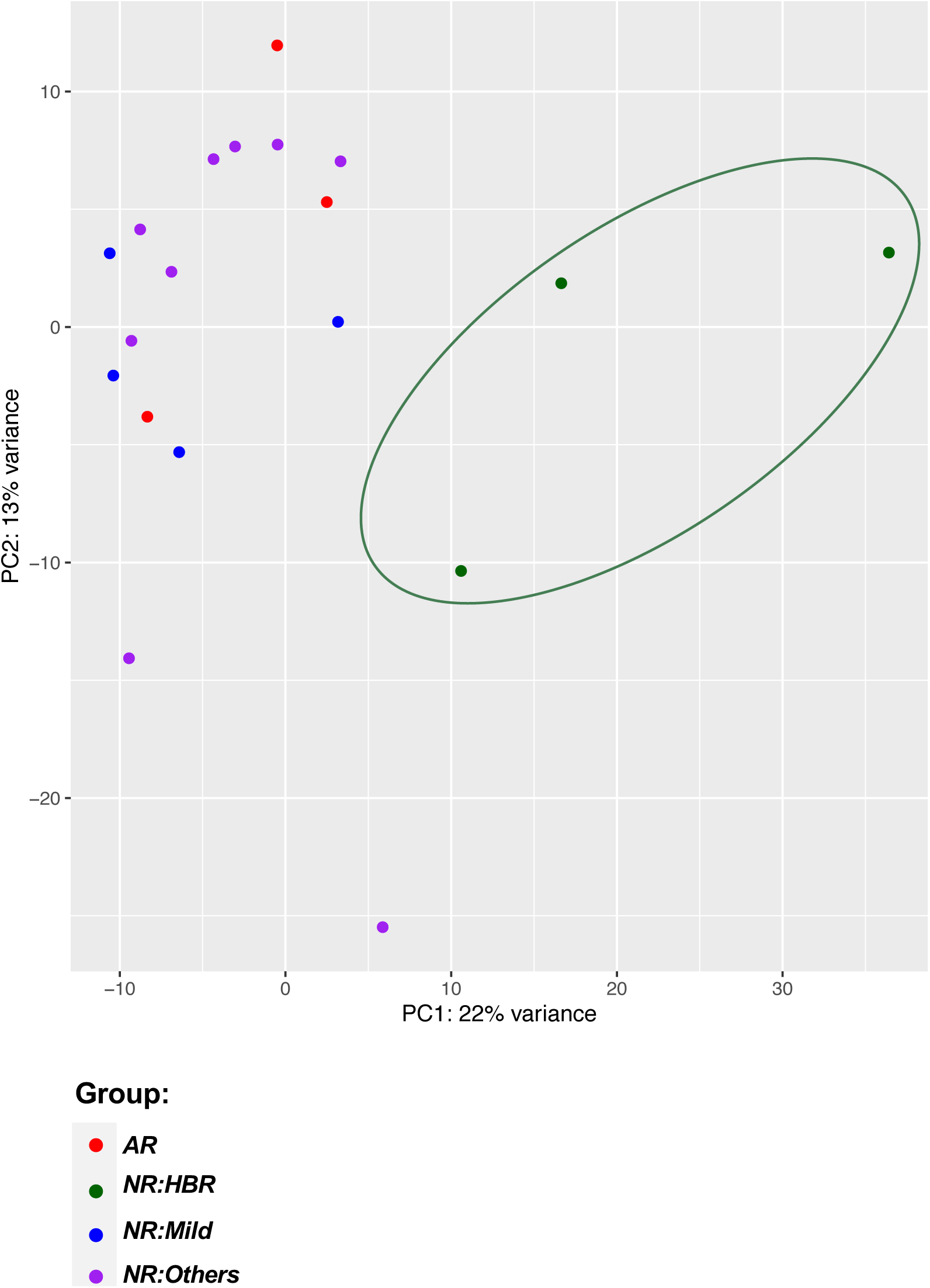
Principal component analysis of RNA-seq from liver biopsies from post-transplantation patients. *AR*, acute rejection; *NR:HBR*, non-rejection with hyperbilirubinemia (serum total bilirubin > 2.5 mg/dL); *NR:Mild*, non-rejection with total bilirubin ≤ 2.5 mg/dL and serum ALT, AST and ALP ≤ 1.67x ULN; *NR:Others*, non-rejection with total bilirubin ≤ 2.5 mg/dL and serum ALT, AST and ALP > 1.67x ULN.

Differential gene expression analysis comparing *NR:HBR* group to all other NR samples showed that 784 genes were differentially expressed. When comparing *NR:HBR* to *NR:Mild*, 977 genes were identified that differentially expressed between the 2 groups. Figure 2A is a volcano plot illustrating the top differentially expressed genes, among which the expression of *CYP7A1* was significantly higher, while *LOXL4, CFTR* and *ADGRG2* expression was lower in the *NR:Mild* group compared to *NR:HBR*. Subsequent GSEA study using the Hallmark pathway database demonstrated increased expression in apoptosis, inflammation and cell proliferation pathways in the *NR:HBR* group compared to *NR:Mild* (Fig. 2B). The Unfolded_Protein_Response pathway is the nineteenth highest enriched pathway, having a normalized enrichment score of 1.431951, although the FDR q-value was 0.051 (Fig 2C). Complementary pathway analysis using GeneCodis revealed that three UPR-related pathways were significantly upregulated in the *NR:HBR* group compared to *NR:Mild* (P-adj<0.05): 1) response to unfolded protein, 2) endoplasmic reticulum unfolded protein response, and 3) negative regulation of PERK-mediated unfolded protein response (Table 2).

**Table 2:**
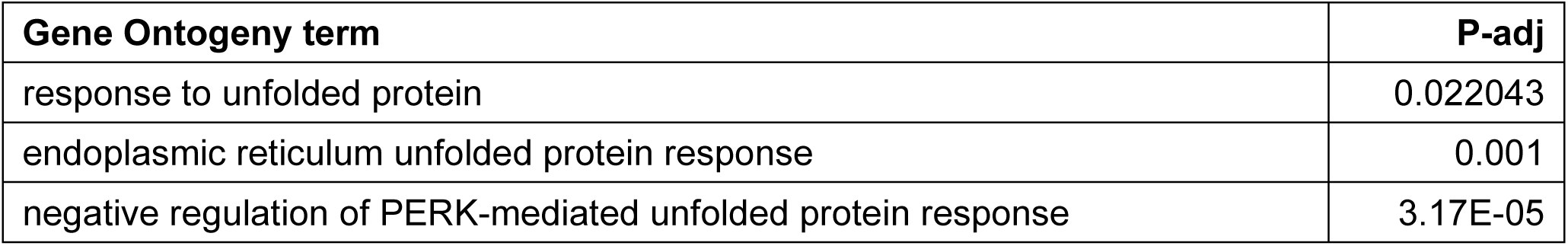
Gene Ontogeny pathway analysis comparing the *NR:HBR* and the *NR:Mild* group using GeneCodis.

**Figure 2:**
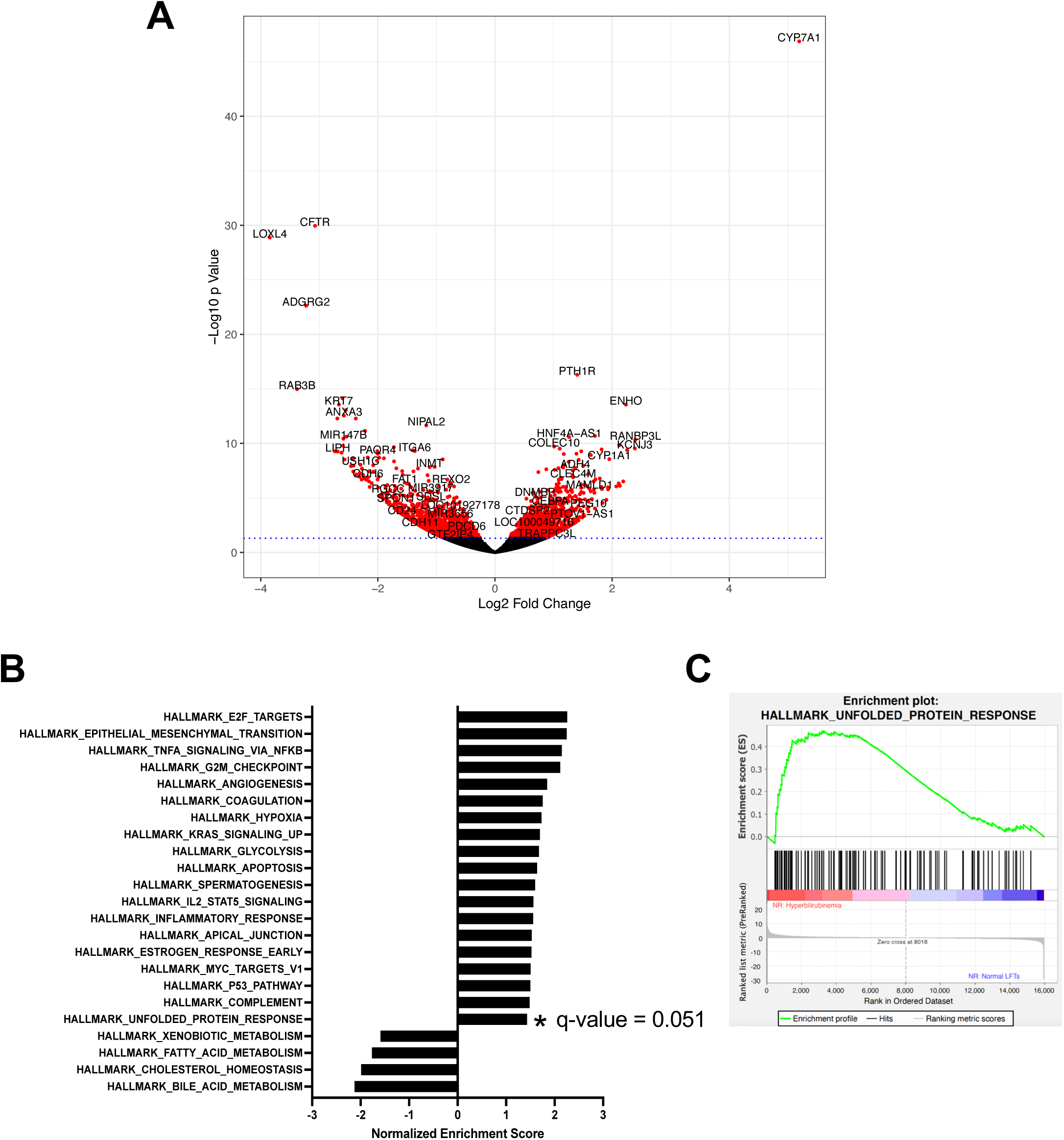
Volcano plot and pathway analysis examining hepatic gene expression in patients with hyperbilirubinemia. (A) Volcano plot comparing the hepatic gene expression of the *NR:HBR* group to the *NR:Mild* group. Expression of *CYP7A1* was significantly higher, while *LOXL4, CFTR* and *ADGRG2* expression was lower in the *NR:Mild* group. (B) Differentially expressed hepatic pathways in patients with hyperbilirubinemia. GSEA study using the Hallmark pathway database comparing the *NR:HBR* group to the *NR:Mild* group. (C) The Unfolded_Protein_Response pathway had a normalized enrichment score of 1.431951 comparing the *NR:HBR* group to *NR:Mild* group, although the FDR q-value was 0.051.

We next compared the RNA-seq expression of liver biopsies from the *AR* group to the *NR* groups. The PCA plot demonstrated that the *AR* group did not cluster independently from the *NR* groups (Figure 1). Lymphocyte activating 3 (*LAG3*) and cyclin dependent kinase 1 (*CDK1*) genes had the greatest increase in expression in *AR* compared to *NR* groups as shown in the volcano plot (Figure 3A). Although several additional genes had changes in gene expression level with a P<0.05, only *LAG3* and *CDK1* had P-adj values less than 0.05. Figure 3B demonstrated that gene expression in the *AR* group was approximately 2.4-fold and 3.4-fold higher, for *LAG3* and *CDK1*, respectively compared to the *NR* groups.

**Figure 3:**
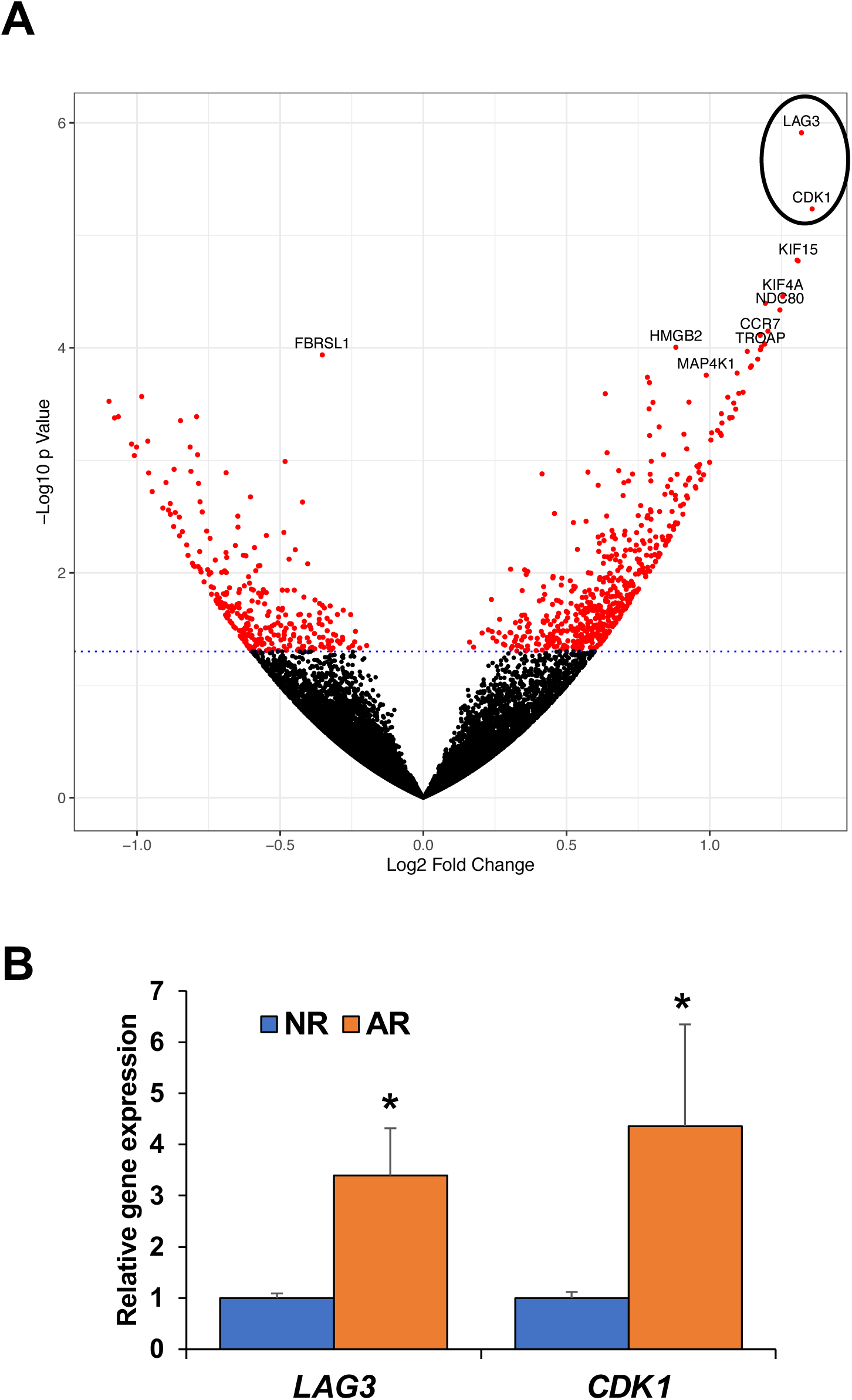
RNA-seq comparing hepatic gene expression in patients with and without acute rejection. (A) Volcano plot demonstrated that the acute rejection (AR) group had increased hepatic gene expression of lymphocyte activating 3 (*LAG3*) and cyclin dependent kinase 1 (*CDK1*) compared to non-rejection (NR) groups. (B) Hepatic gene expression of *LAG3* and *CDK1* in patients with AR and NR. *P-adj < 0.05.

We subsequently sought to determine, using all samples, if the level of the serum liver chemistries (ALT, AST, total bilirubin and ALP) correlated with the expression level of hepatic UPR genes. Table 3 lists the Pearson r and P-values of the Pearson Correlation analysis comparing levels of serum ALT, AST and total bilirubin, with expression of the UPR genes from the XBP1, PERK and ATF6 pathways. Supplementary Figure S1-S3 are the graphs of these data. Overall, one of the PERK pathway target genes, activating transcription factor 3 (*ATF3*) showed the highest correlations with aminotransferase levels (r = 0.77 with p = 0.001 for ALT, and r = 0.68 with p = 0.001 for AST), whereas *ATF6* gene expression correlated most with total bilirubin (r = 0.69 with p = 0.001). Of note, no correlations were identified between serum ALP levels and the expression of liver UPR genes. Tables 4 and 5 list the Pearson r and P-values comparing serum ALT, AST, ALP and total bilirubin to the expression of bile acid metabolism and inflammatory genes. Supplementary Figures S4 and S5 are the graphs of these data. Among genes tested, fibroblast growth factor 19 (*FGF19)* known to play a key role in regulating bile acid synthesis had the highest correlation with total bilirubin with r of 0.96 (p = 0.0001). Two bile acid transporters genes, *SLC51B* encoding organic solute transporter beta and *ABCB4* encoding ATP binding cassette subfamily B member 4 (also known as PFIC-3), correlated with all 4 serum liver chemistries. Inflammatory gene *FOXP3* expression had the highest correlation with total bilirubin (r = 0.8283, p = 0.0001) and it also correlated with ALT and AST.

**Table 3:** Pearson correlation analysis of serum liver chemistries and hepatic UPR gene expression.

| Unfolded Protein Response Pathway | Gene | Pearson r | p-value |
| --- | --- | --- | --- |
| **Correlated with ALT** |  |  |  |
| XBP1 pathway | *FICD* | 0.5545 | 0.0138 |
| PERK pathway | *ATF3* | 0.773 | 0.0001 |
| ATF6 pathway | *MIS12* | 0.5015 | 0.0287 |
|  | *DNAJB11* | 0.4854 | 0.0351 |
| **Correlated with AST** |  |  |  |
| XBP1 pathway | *FICD* | 0.6689 | 0.0017 |
|  | *STT3A* | 0.5103 | 0.0256 |
|  | *MBNL2* | 0.4659 | 0.0444 |
|  | *HSPA13* | 0.681 | 0.0013 |
|  | *SEC23B* | 0.6366 | 0.0034 |
| PERK pathway | *WARS* | 0.5072 | 0.0267 |
|  | *ATF3* | 0.6847 | 0.0012 |
|  | *PPP1R15A* | 0.5843 | 0.0086 |
| ATF6 pathway | *HYOU1* | 0.5871 | 0.0082 |
|  | *MANF* | 0.6064 | 0.0059 |
|  | *HSPA5* | 0.5272 | 0.0204 |
|  | *PDIA6* | 0.5355 | 0.0181 |
|  | *DNAJB11* | 0.6268 | 0.0041 |
| **Correlated with total bilirubin** |  |  |  |
| PERK pathway | *PPP1R15A* | 0.6351 | 0.0035 |
|  | *PON2* | 0.5278 | 0.0202 |
|  | *ATF6* | 0.6891 | 0.0011 |
| ATF6 pathway | *HYOU1* | 0.5111 | 0.0253 |
|  | *MANF* | 0.516 | 0.0237 |
|  | *HSPA5* | 0.5366 | 0.0178 |
|  | *HERPUD1* | 0.4576 | 0.0488 |
|  | *SEL1L* | 0.5285 | 0.02 |
|  | *DNAJB11* | 0.6193 | 0.0047 |

**Table 4:** Pearson correlation analysis of serum liver chemistries and hepatic bile acid metabolism gene expression.

| **Bile acid metabolism** |  |  |
| --- | --- | --- |
| **Gene** | **Pearson r** | **p-value** |
| **Correlated with ALT** |  |  |
| *SLC51B* | 0.6562 | 0.0023 |
| *ABCB4* | 0.6548 | 0.0035 |
| *SLCO1B1* | -0.5436 | 0.016 |
| *FGF19* | 0.6408 | 0.0031 |
| **Correlated with AST** |  |  |
| *SLC51B* | 0.5846 | 0.0086 |
| *ABCB4* | 0.5729 | 0.0104 |
| *SLCO1B1* | -0.5458 | 0.0156 |
| *FGF19* | 0.5148 | 0.0241 |
| **Correlated with ALP** |  |  |
| *SLC51B* | 0.4799 | 0.0376 |
| *ABCB4* | 0.5087 | 0.0262 |
| **Correlated with total bilirubin** |  |  |
| *SLC51A* | 0.6292 | 0.0039 |
| *SLC51B* | 0.7521 | 0.0002 |
| *ABCB4* | 0.5474 | 0.0153 |
| *FGF19* | 0.9583 | 0.0001 |

**Table 5:** Pearson correlation analysis of serum liver chemistries and hepatic inflammation gene expression.

| **Inflammation** |  |  |
| --- | --- | --- |
| **Gene** | **Pearson r** | **p-value** |
| **Correlated with ALT** |  |  |
| *CD163* | 0.5149 | 0.0241 |
| *FOXP3* | 0.6089 | 0.0057 |
| *CCL2* | 0.4971 | 0.0304 |
| **Correlated with AST** |  |  |
| *IFNG* | 0.4563 | 0.0495 |
| *CD163* | 0.6388 | 0.0032 |
| *ICAM1* | 0.5384 | 0.0174 |
| *FOXP3* | 0.5474 | 0.0153 |
| *CCL2* | 0.4888 | 0.0337 |
| *CD3D* | 0.4624 | 0.0462 |
| *CD8A* | 0.4831 | 0.0362 |
| **Correlated with total bilirubin** |  |  |
| *FOXP3* | 0.8283 | 0.0001 |
| *CCL2* | 0.5469 | 0.0154 |

## Discussion

In this study, we performed hepatic transcriptome analysis on human liver biopsies from post-liver transplantation patients. Pathway analysis demonstrated increased expression of the liver UPR pathways in the *NR:HBR* group compared to the *NR:Mild* group. It has been previously reported that selected UPR genes are down-regulated in PSC patients with a high risk of developing PSC-related complications (7). Patients with progressive NASH also have UPR dysregulation and an attenuated UPR response compared to patients with benign hepatic steatosis (24), although this was not seen in other patient populations (25). Similarly, weanling mice have an impaired ability to activate their hepatic XBP1 pathway, with a resultant increase in serum ALT, increased proapoptotic C/EBP homologous protein and death receptor 5 expression, and enhanced liver apoptosis (26). Therefore, hepatic UPR activation may be a protective response to cholestatic liver injury, while an impaired or attenuated UPR response can lead to increased liver injury in both animal models of cholestasis and potentially human diseases such as PSC and NASH.

We next sought to determine the relationship between the levels of serum liver chemistries and expression of hepatic UPR genes. We identified significant correlations between serum ALT and AST with downstream gene targets of all three UPR pathways (XBP1, PERK and ATF6), and total bilirubin correlated with downstream targets of PERK and ATF6 pathways. Although these correlations do not imply a causative relationship, it is worth noting that there was a consistently positive relationship between increasing levels of these serum liver chemistries and increasing UPR gene expression. This finding further supports the relationship between increased ER stress and degree of hepatocellular injury and/or diminished hepatobiliary secretory function.

Although comparing the *AR* group with the *NR* groups did not reveal independent gene clustering or altered gene expression of UPR pathways, the *AR* group had significantly increased gene expression of *LAG3* and *CDK1*. LAG3 is highly expressed in activated T-lymphocytes, and the increased expression that we observed using bulk RNA-seq may be due to intrahepatic T-lymphocyte activation rather than enhanced expression in primary liver parenchyma. Single cell or single-nuclei RNA-seq provides more in-depth information on cell-specific gene expression, however our samples were stored in RNAlater and not suitable for such experiments. In addition, the increased expression of *CDK1*, a key gene in cell cycle, in the *AR* group is likely due to enhanced cell proliferation that can occur in response to hepatic injury. There were a relatively small number of patients with acute rejection, and it is possible that other associations can be identified using a larger patient population (27-29).

We defined our cholestasis patient group using serum total bilirubin rather than serum bile acid levels since serum bile acid levels were not routinely obtained in our patient groups. It is well accepted that one of the major factors causing cholestatic liver injury is increased hepatocellular bile acid concentrations, hydrophobicity and/or a total bile acid pool. Animal studies using bile acid toxicity models have demonstrated that cholestasis induces ER stress and UPR activation (8, 26). Of note, induction of hepatic ER stress in cholestasis decreases gene expression of *Cyp7a1, Fxr, Abcc3* and *Abcb11* similar to the pattern observed in our study (30, 31). These hepatic changes can reduce bile acid synthesis and increase bile acid efflux transporters, which are protective responses to reduced hepatocellular bile acid toxicity. There are additional causes of serum bilirubin elevations including increased bilirubin formation from severe internal bleeding, multiple blood transfusions, hemolysis or dyserythropoiesis. However, there was no evidence for these alternate etiologies in our patient cohort.

The liver biopsy specimens utilized for the study were from a patient population with previous liver transplantation, obtained for-cause but otherwise in an unbiased manner. It is possible that immunosuppressive and other medications could potentially affect hepatic gene expression. Therefore, it would be interesting to extend these observations using liver biopsies from other patient populations. A growing literature of murine data has demonstrated the causative relationship between cholestasis, and ER stress with UPR activation, with a paucity of data in human populations. Our liver biopsy transcriptome data provide a novel demonstration of an association between human hepatic UPR gene expression and human cholestasis.

## Acknowledgement

The authors wish to thank the Northwestern University Center of Genetic Medicine NUSeq core facility for performing RNA-seq and for assistance in bioinformatics analysis.

**Author names in bold designate shared co-first authorship**.

