## Supplementary Table 1 for "Expression of Unfolded Protein Response Genes in Post-transplantation Liver Biopsies"

**Supplementary Table 1: Individual patient characteristics.**

| Groups | Sex | Age (y) | Race | Indication for OLT | ALT (U/L) | AST (U/L) | ALP (U/L) | Total Bilirubin (mg/dL) |
|---|---|---|---|---|---|---|---|---|
| *AR* | M | 47 | White | Alcohol, A-1AD | 252 | 141 | 226 | 1.2 |
| *AR* | F | 20 | White | Wilson Disease | 68 | 27 | 54 | 0.5 |
| *AR* | F | 64 | White | NASH | 195 | 178 | 157 | 1 |
| *NR:HBR* | M | 27 | White | PSC/AIH | 302 | 127 | 366 | 12.2 |
| *NR:HBR* | F | 43 | White | Primary Sclerosing Cholangitis | 27 | 50 | 164 | 3.5 |
| *NR:HBR* | M | 65 | White | Primary Biliary Cholangitis | 165 | 134 | 261 | 7.4 |
| *NR:Mild* | M | 26 | Other | Primary Sclerosing Cholangitis | 31 | 35 | 130 | 0.5 |
| *NR:Mild* | M | 65 | White | Alcoholic | 20 | 18 | 87 | 1.5 |
| *NR:Mild* | M | 35 | White | Autoimmune | 84 | 48 | 68 | 0.7 |
| *NR:Mild* | F | 62 | White | Cryptogenic | 79 | 64 | 113 | 0.6 |
| *NR:Others* | F | 65 | Hispanic/Latino | Cryptogenic, HCC | 303 | 103 | 432 | 1.5 |
| *NR:Others* | F | 58 | White | NASH | 22 | 24 | 303 | 0.6 |
| *NR:Others* | M | 25 | White | Biliary Atresia | 75 | 47 | 621 | 1.1 |
| *NR:Others* | F | 65 | White | Primary Biliary Cholangitis | 24 | 24 | 235 | 0.5 |
| *NR:Others* | F | 62 | White | Primary Biliary Cholangitis, HCC | 152 | 22 | 221 | 1.3 |
| *NR:Others* | M | 45 | White | Primary Sclerosing Cholangitis | 39 | 37 | 195 | 0.9 |
| *NR:Others* | F | 59 | White | Primary Biliary Cholangitis | 41 | 29 | 533 | 0.3 |
| *NR:Others* | M | 62 | Back | HCV | 105 | 40 | 191 | 1.2 |
| *NR:Others* | M | 43 | White | Cryptogenic | 124 | 74 | 532 | 1.5 |

***AR***: acute rejection

***NR:HBR***: non-rejection with hyperbilirubinemia (serum total bilirubin > 2.5 mg/dL)

***NR:Mild***: non-rejection; serum total bilirubin ≤ 2.5 mg/dL; ALT, AST and ALP ≤ 1.67x ULN

***NR:Others***: other non-rejection with serum total bilirubin ≤ 2.5 mg/dL; ALT, AST and ALP > 1.67x ULN
