## Supplementary Figure legends for "Expression of Unfolded Protein Response Genes in Post-transplantation Liver Biopsies"

**Supplementary Figure S1: Hepatic unfolded protein response gene expression correlated with serum ALT.** Graphs demonstrating the Pearson correlation between the serum ALT levels and hepatic gene expression of the downstream targets of the XBP1, PERK and ATF6 pathways.

**Supplementary Figure S2: Hepatic unfolded protein response gene expression correlated with serum AST.** Graphs demonstrating the Pearson correlation between the serum AST levels and hepatic gene expression of the downstream targets of the XBP1, PERK and ATF6 pathways.

**Supplementary Figure S3: Hepatic unfolded protein response gene expression correlated with serum total bilirubin.** Graphs demonstrating the Pearson correlation between the serum total bilirubin levels and hepatic gene expression of the downstream targets of the PERK and ATF6 pathways.

**Supplementary Figure S4: Hepatic bile acid metabolism gene expression correlated with serum liver chemistries.** Pearson correlation graphs demonstrated the hepatic expression of several bile acid metabolism genes correlated with serum ALT, AST, ALP and/or total bilirubin.

**Supplementary Figure S5: Hepatic inflammation gene expression correlated with serum liver chemistries.** Pearson correlation graphs demonstrated the hepatic expression of several inflammatory genes correlated with serum ALT, AST and/or total bilirubin.
