## Supplementary figures and images for "Expression of Unfolded Protein Response Genes in Post-transplantation Liver Biopsies"

### Supplementary Figure S1

#### Supplementary Figure S1

##### XBP1 pathway

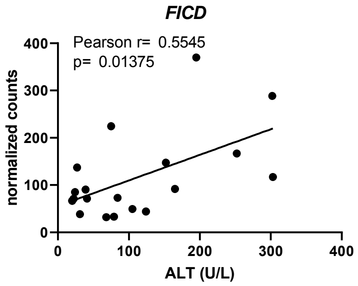


##### PERK pathway

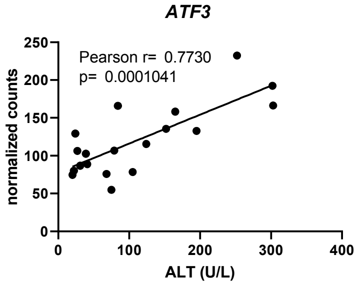


##### ATF6 pathway

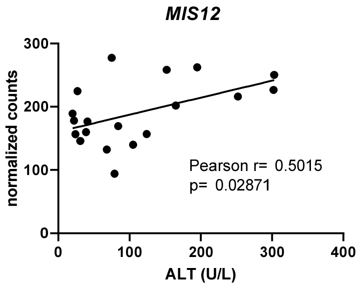


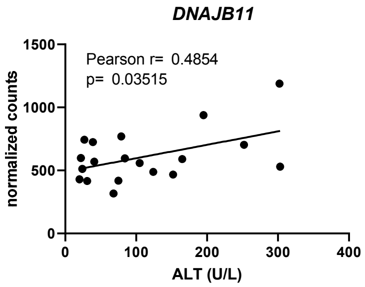

### Supplementary Figure S2

#### Supplementary Figure S2

##### XBP1 pathway

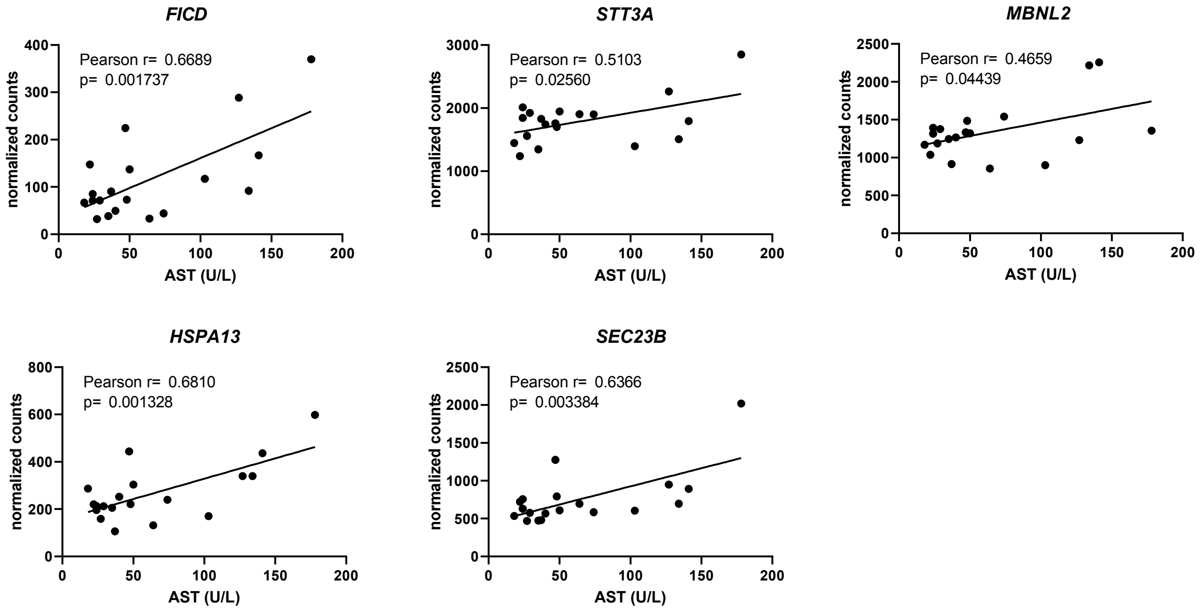


##### PERK pathway

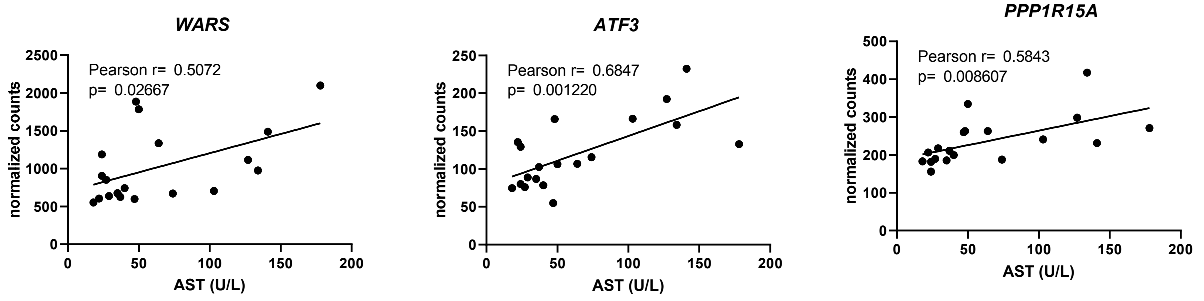


##### ATF6 pathway

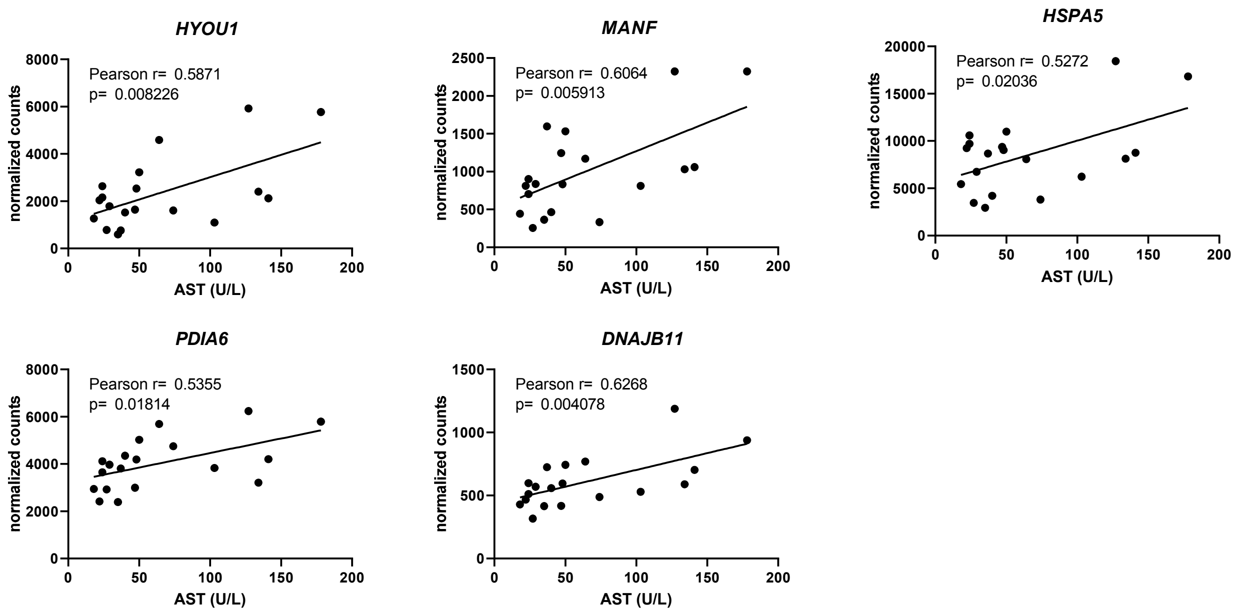

### Supplementary Figure S3

#### Supplementary Figure S3

##### PERK pathway

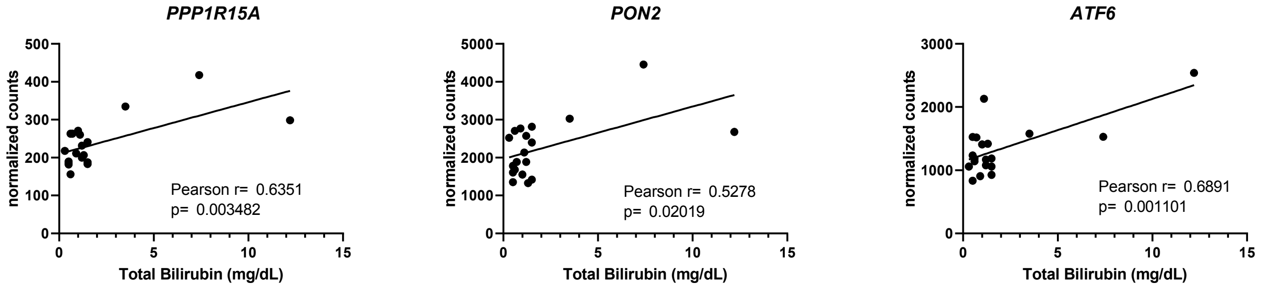


##### ATF6 pathway

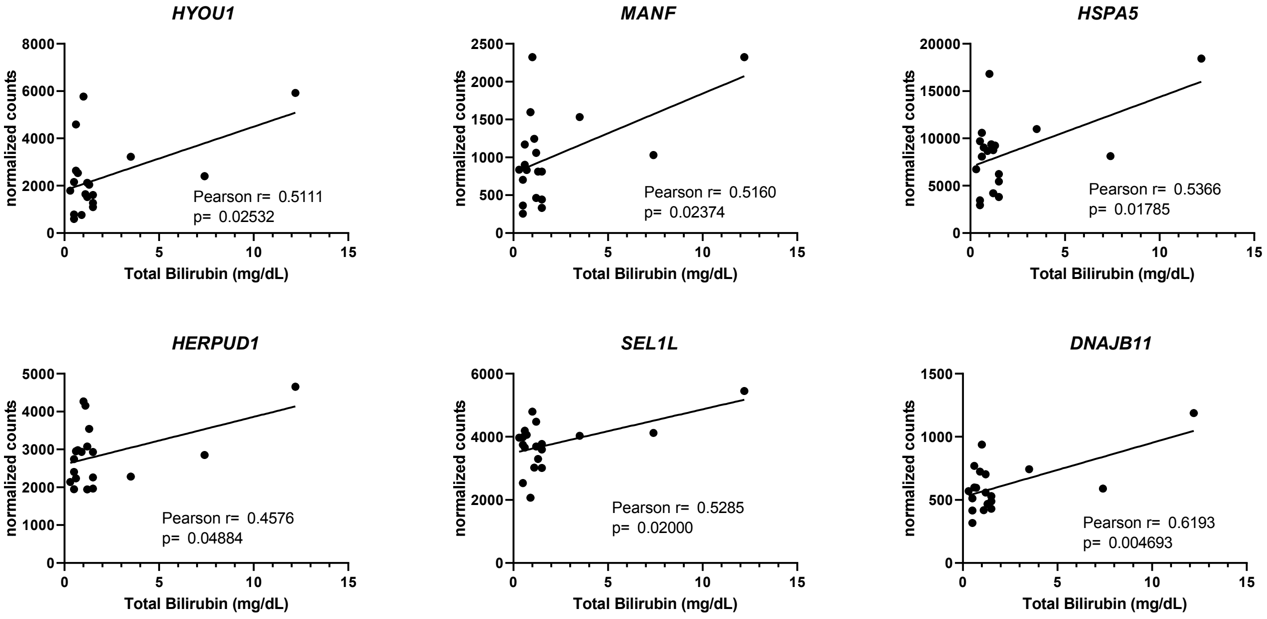
