## Supplementary Figure S5 for "Expression of Unfolded Protein Response Genes in Post-transplantation Liver Biopsies"

### Correlated with ALT

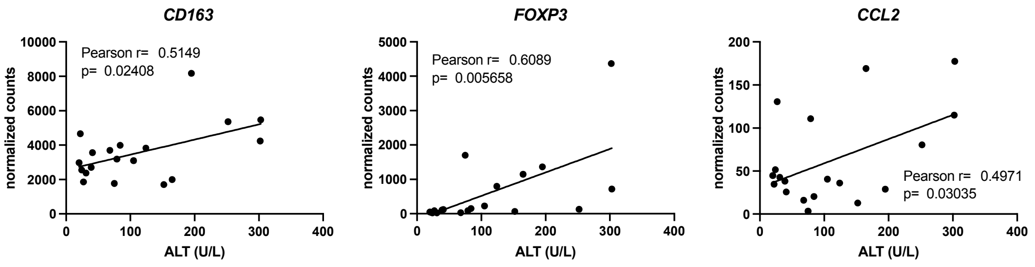


### Correlated with AST

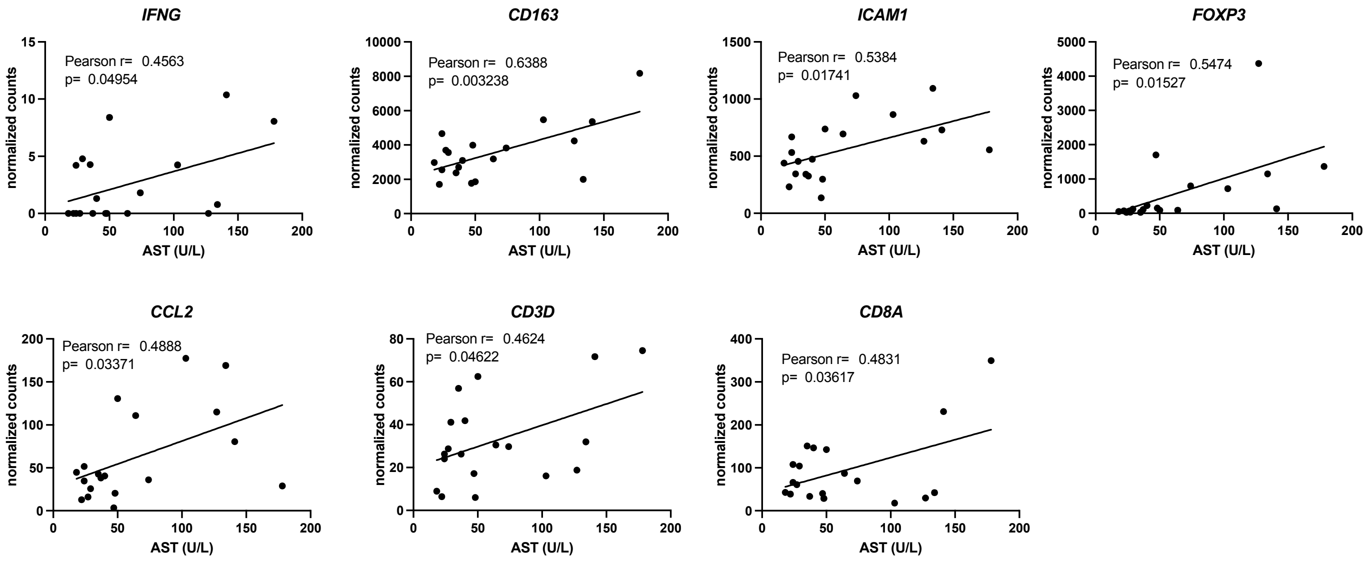


### Correlated with Total Bilirubin

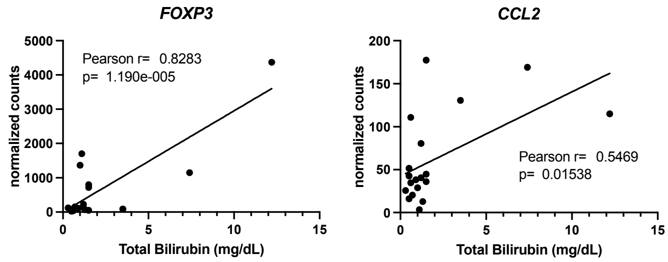
